# Acidogenic Fermentation of Sugarcane Vinasse and Glucose Using Two Different Anaerobic Digestion Inocula

**DOI:** 10.1101/2021.02.22.432247

**Authors:** Juliana Martins, Hugo Valença de Araújo, Gustavo Mockaitis, Ariovaldo José da Silva

**Affiliations:** School of Agricultural Engineering (FEAGRI), University of Campinas (UNICAMP), Av. Cândido Rondon, 501, Barão Geraldo, Campinas, São Paulo 13083-875, Brazil; Technology Center (CTEC), Federal University of Alagoas (UFAL), Campus A. C. Simões, Av. Lourival Melo Mota, Tabuleiro dos Martins, Maceió, Alagoas 57072-970, Brazil

**Keywords:** anaerobic digestion, batch reactors, lactic acid, ethanol, kinetic modelling

## Abstract

Vinasse is an agroindustrial wastewater obtained at large amounts in countries such as Brazil, which is one of the world greatest sugar and ethanol producers from sugarcane. Despite its most common use is as nutrient for fertirrigation in sugarcane crops nowadays, alternative processing has been intensely sought in last decades for vinasse aiming primarily the production of energy carriers and high-added value chemicals. One of these alternatives is the anaerobic digestion promoted by bacteria, which results in several products of economic importance. In view of these information, this work presents an investigation of acidogenic fermentation by *C. acetobutylicum* ATCC 824 and *C. beijerinckii* ATCC 25752 was conducted in batch reactors by using synthetic vinasse and aqueous glucose solution as substrates. Evaluation of effects of microorganism and carbon source was focused mainly on the production of short chain organic acids (SCOAs), although other metabolites, such as ethanol, were also detected. Carbohydrates were effectively consumed by bacteria, so remaining concentrations of sucrose and glucose (Suc and Glu) in both substrates were considerably low at the end of tests. Lactic acid (HLa) was the prevailing metabolite in all experiments, determining the trends observed for total SCOAs content as a function of incubation time, while other acids were generated in much lesser amounts. Moreover, ethanol (EtOH) attained notably higher concentrations in batch reactors containing glucose as substrate, which showed more relevant production of this chemical. Maximum of 14.1 g HLa L^−1^ was quantified for lactic acid at incubation time 96 h in reactor using synthetic vinasse and *C. beijerinckii* (VB), linked to a productivity *P* = 146.46 mg HLa L^−1^ h^−1^ and a yield *Y* = 1108.03 mg HLa (g Sue)^−1^. For ethanol, maximum was 1.6 g EtOH L^−1^, occurring in reactor containing glucose and *C. acetobutylicum* (GA) also at incubation time 90 h, resulting in *P* = 13.60 mg EtOH L^−1^ h^−1^ and *Y* = 73.04 mg EtOH (g Glu)^−1^. Finally, kinetic models were adjusted to experimental data of carbohydrate content, lactate concentration and optical density (indicative of microorganism growth), showing good prediction capability as suggested by values of fit quality parameters such as Residual Sum of Squares (*RSS*) and *R*^2^.

## 1. INTRODUCTION

More than ever, one of the main concerns of research in energy and sanitation engineering nowadays is on how to avoid impacts due to disposition of wastewaters in environment and also on how to use residuals to generate added-value products such as energy carriers and chemicals with applications in relevant areas. Within this scope, acidogenic fermentation of liquid wastes from different origins (industrial, domestic or even the synthetic ones) is an alternative that has been studied intensively. Acidogenesis is a biological process and depending on which microorganism is used, its major products are some short chain organic acids (SCOAs) and high-energy content biogas such as hydrogen and methane (MORAES *et al.*, 2019; WAINAINA *et al.*, 2019). Thus, renewability of daily life and also of large scale industrial activity can be certainly increased by means of using acidogenic fermentation to convert obtained liquid residuals into useful energy and material inputs for society.

In the Brazilian context, industry of sugar and ethanol from sugarcane presents great economic importance since both products provide much of internal food, beverage and fuel markets besides also account for a large portion of country exports (WILKINSON, 2015). Productive process of sugar and ethanol results in two main residues, namely sugarcane bagasse, obtained right after extraction of sugarcane juice, and sugarcane vinasse, generated as bottom stream from hydrous ethanol distillation column. Sugarcane bagasse has been used for a long time as a solid fuel for burning in boilers existing on sugar and ethanol mills aiming to generate steam and electricity to supply internal and even external requirements (ENSINAS *et al.*, 2007). Vinasse has been traditionally applied as a nutrient for fertigation of sugarcane crops because of its high contents of nitrogen, phosphorus, potassium (NPK) and other essential mineral elements for plant development (JIANG *et al.*, 2012). However, seeing that continued use of vinasse on soil can result in a number of harmful environmental consequences (FUESS & GARCIA, 2014), various studies carried out during last decades have focused on unconventional ways of processing it mainly to obtain high added-value products (HOARAU *et al.*, 2018).

It is estimated that sugar and ethanol mills produce an average ratio of 13 L of vinasse for each liter of ethanol (GUNKEL *et al.*, 2006). However, it can vary widely (FREIRE e CORTEZ, 2000; CAVALLET *et al.*, 2012) as a consequence of factors such as feedstock quality, particular conditions of industrial unit operations, material used for must preparation and yield of ethanol fermentation process (WILKIE *et al.*, 2000; PANT & ADHOLEYA, 2007). Therefore, based on typical average ratios, reasonable projections have indicated a generation of 1,742 million of liters of vinasse as agroindustrial wastewater since 2018 until 2024 (HOARAU *et al.*, 2018). In view that this amount is very significant, it not solely justifies, but also motivates studies on novel technologies capable of converting sugarcane vinasse into biofuels and valuable chemicals (MORAES *et al.*, 2019; FUESS *et al.*, 2020).

Anaerobic digestion of vinasse has been orientated mostly to generation of hydrogen and methane (biogas for use as biofuel), so there is a lack of works regarding use of vinasse as substrate aiming the production of SCOAs and solvents (MORAES *et al.*, 2019), which serves as an incentive for further exploration on this topic. Acidogenesis products such as butyric, acetic and propionic acids have found uses in textile, pharmaceutical, food, beverage, fragrance, cosmetics, pesticides and some other industrial sectors (DWIDAR *et al.*, 2012; HIRAL *et al.*, 2017). Among all of them, lactic acid (lactate) is certainly one of the most wanted acidic metabolites due to its great versatility, which makes it useful for applications as acidifying and flavouring agent, leather softener, antimicrobial preservative and monomer in commercial chemistry (HIRAL *et al.*, 2017; HOARAU *et al.*, 2018).

In this work, main goal was to investigate the acidogenic fermentation performed by *C. acetobulycum* ATCC 824 and *C. beijerinckii* ATCC 25752 strains on two wastewaters prepared at laboratory, namely aqueous glucose solution and a synthetic vinasse, with controlled characteristics. Both bacteria are classically employed in solvent production, specially acetone, butanol and ethanol (ABE fermentation), however it is well known that first phase of their metabolic routes are composed by chemical reactions providing acidic products (KUMAR & GAYEN, 2011; LÜTKE-EVERSLOH & BAHL, 2011). Synthetic vinasse was aimed to simulate the industrial liquid residue, but avoiding the great variability typical of it, so it is possible to verify the best conditions for fermentation process more easily, even if at a preliminary stage. Therefore, it is believed that major contributions given here concerns obtaining results of batch experiments of acidogenesis with the selected microbes and substrates and fitting kinetic models to some monitoring parameters of process.

## 2. MATERIALS & METHODS

### 2.1 Substrates

One of the substrates studied in this work was a synthetic vinasse prepared in the Laboratory of Environment and Sanitation (LMAS) of School of Agricultural Engineering of University of Campinas (FEAGRI/UNICAMP). It presents similar characteristics to those of soluble fraction of real sugarcane vinasse aiming to simulate as closest as possible the wastewater from sugar and ethanol industry. Composition of a synthetic vinasse proposed by Godoi *et al.* (2017) was taken as a reference in order to define composition of synthetic vinasse used in batch reactors of the present work. Thus, composition applied here is depicted in Table 1 for a liquid residue at 20 g COD L^−1^, which was the value adopted for soluble chemical oxygen demand (CODsol).

**Table 1.** Composition of synthetic vinasse used in the present work.

| Constituent | Concentration | Reactant | Solution | Synthetic vinasse<sup>+</sup> |
| --- | --- | --- | --- | --- |
| COD<sub>sol</sub> (g O<sub>2</sub> L<sup>-1</sup>) | 20.0 | – | – | – |
| Propionic acid (g HPr L<sup>-1</sup>) | 0.50 | C<sub>3</sub>H<sub>6</sub>O<sub>2</sub> | P.A (99.9%) | 0.51 mL L<sup>-1</sup> |
| Butyric acid (g HBu L<sup>-1</sup>) | 0.70 | C<sub>4</sub>H<sub>8</sub>O<sub>2</sub> | P.A (99.7%) | 0.73 mL L<sup>-1</sup> |
| Acetic acid (g HAc L<sup>-1</sup>) | 1.30 | C<sub>2</sub>H<sub>4</sub>O<sub>2</sub> | P.A (99.7%) | 1.24 mL L<sup>-1</sup> |
| Ethanol (g EtOH L<sup>-1</sup>) | 1.50 | C<sub>2</sub>H<sub>6</sub>O | P.A (95.0 %) | 2.00 mL L<sup>-1</sup> |
| Phenol (mg PhOH L<sup>-1</sup>) | 6.0 | C<sub>6</sub>H<sub>5</sub>OH | 12 g L<sup>-1</sup> | 0.50 mL L<sup>-1</sup> |
| Phosphorus (mg P-PO<sub>4</sub><sup>3+</sup> L<sup>-1</sup>) | 70.0 | K<sub>2</sub>HPO<sub>4</sub> | 78.734 g L<sup>-1</sup> | 2.50 mL L<sup>-1</sup> |
|  |  | KH<sub>2</sub>PO<sub>4</sub> | 61.510 g L<sup>-1</sup> |  |
| Iron (mg Fe L<sup>-1</sup>) | 10.0 | FeCl<sub>3</sub>.6H<sub>2</sub>O | 96.759 g L<sup>-1</sup> | 0.50 mL L<sup>-1</sup> |
| Manganese (mg Mn L<sup>-1</sup>) | 3.00 | MnSO<sub>4</sub>.H<sub>2</sub>O | 18.452 g L<sup>-1</sup> |  |
| Zinc (mg Zn L<sup>-1</sup>) | 0.40 | ZnCl<sub>2</sub> | 2.668 g L<sup>-1</sup> |  |
| Copper (mg Cu L<sup>-1</sup>) | 0.30 | CuCl<sub>2</sub>.2H<sub>2</sub>O | 1.609 g L<sup>-1</sup> | 0.50 mL L<sup>-1</sup> |
| Nickel (mg Ni L<sup>-1</sup>) | 0.20 | NiCl<sub>2</sub>.6H<sub>2</sub>O | 1.608 g L<sup>-1</sup> |  |
| Lead (mg Pb L<sup>-1</sup>) | 0.20 | Pb(NO<sub>3</sub>)<sub>2</sub> | 0.639 g L<sup>-1</sup> | 0.50 mL L<sup>-1</sup> |
| Cadmium (mg Cd L<sup>-1</sup>) | 0.08 | Cd(NO<sub>3</sub>)<sub>2</sub>.4H<sub>2</sub>O | 0.439 g L<sup>-1</sup> |  |
| Sucrose (g Suc L<sup>-1</sup>) | 12.00 | Raw sugar | weighted* | 12.50 g L<sup>-1</sup> |
| Nitrogen (g N L<sup>-1</sup>) | 0.80 | CH<sub>4</sub>N<sub>2</sub>O | weighted* | 1.71 g L<sup>-1</sup> |
| Potassium (g K L<sup>-1</sup>) | 2.20 | K<sub>2</sub>SO<sub>4</sub> | weighted* | 1.81 g L<sup>-1</sup> |
|  |  | KCl | weighted* | 2.23 g L<sup>-1</sup> |
| Calcium (g Ca L<sup>-1</sup>) | 0.44 | CaCl<sub>2</sub>.2H<sub>2</sub>O | weighted* | 1.61 g L<sup>-1</sup> |
| Magnesium (g Mg L<sup>-1</sup>) | 0.25 | MgCl<sub>2</sub>.6H<sub>2</sub>O | weighted* | 2.09 g L<sup>-1</sup> |
| Sodium (g Na L<sup>-1</sup>) | 0.52 | NaCl | weighted* | 1.32 g L<sup>-1</sup> |
| Sulphate (g S-SO<sub>4</sub><sup>2-</sup> L<sup>-1</sup>) | 1.00 | – | – | – |

\*Weighted for each liter of synthetic vinasse; <sup>+</sup> Amounts required for preparing 1 L of synthetic vinasse at 20 g COD L<sup>-1</sup>.

Amounts required for preparing synthetic vinasse at 20 g COD L^−1^ were calculated by means of stoichiometry of oxidation reactions of constituents. Sulphate concentration resulted from summation of all reactants containing SO_4_^2-^ anion, namely MnSO_4_.H_2_O and K_2_SO_4_. Potassium concentration was obtained directly by reactants K_2_SO_4_ and KCl, but also indirectly by K_2_HPO_4_ and KH_2_PO_4_, both used to supply phosphorus. Moreover, it is important to highlight that some groups of chemical elements were provided by the same initial solution, such as iron, manganese and zinc; copper and nickel; lead and cadmium.

Additionally, aqueous solution of glucose was also studied as substrate. It was prepared by simple dissolution of primary standard (P.A) solid glucose (Glu) in distilled water, resulting in a solution with a concentration of approximately 20 g Glu L^−1^ (near to 20 g COD L^−1^ of CODsol for synthetic vinasse).

### 2.2 Inoculum

Pure cultures of *Clostridium acetobutylicum* ATCC 824 and *Clostridium beijerinckii* ATCC 25752 bacteria strains were used as inocula in this work. They were obtained in ampoules in the lyophilized form from André Tosello Foundation (Campinas/SP, Brazil), both coming from a microorganisms bank. Ampoules were opened inside a pre-sterilized laminar flow chamber and content of each one was transferred to different test tubes to be mixed to 5 mL of Clostridium Nutrient Medium (CNM Sigma-Aldrich^®^) containing the following concentrations of its compounds: 0.5 g L^−1^ agar; 0.5 g L^−1^ L-cysteine hydrochloride; 5.0 g L^−1^ D(+) glucose; 10.0 g L^−1^ meat extract; 5.0 g L^−1^ peptone; 3.0 g L^−1^ sodium acetate; 5.0 g L^−1^ sodium chloride; 1.0 g L^−1^ starch; 3.0 g L^−1^ yeast extract.

Growth of *Clostridium* cultures were executed before they were used in experiments. For accomplishing this, 1 L of CNM was prepared for mixing with each one of bacteria strain. Both media (CNM + bacteria strain) were autoclaved at 121 °C for 15 min. After they were spontaneously cooled down to environment temperature, media were put into Erlenmeyer flasks of 2 L inside the laminar flow chamber pre-sterilized with 70% v/v ethyl alcohol and under UV radiation for 20 min. Each one of Erlenmeyer flasks was filled with 200 L of *C. acetobutylicum* and *C. beijerinckii* pure cultures. Finally, flasks were closed with a sterile blanket and kept in an oven at 35 °C until turbidity of both media was stabilized.

Growing curves of bacteria cultures were monitored via optical density (OD, indicative of medium turbidity), which was measured by means of a spectrophotometer under wavelength λ = 600 nm for absorbance and also by means of Total Volatile Solids (TVS) quantification. Analysis of OD consisted of reading absorbance of samples diluted with distilled water in proportion 1:5 (BEGOT *et al.*, 1996). Absorbance increasing reflects increase of sample turbidity and therefore points out there was bacteria growth (cell multiplication).

### 2.3 Batch acidogenesis experiments

Experiments were performed in batch reactors in duplicate runnings, under 37 °C and continuous agitation in an orbital shaker working at a rotational speed of 100 rpm. Reactors were prepared in Duran® flasks of 1 L occupied by 850 mL of reaction volume (150 mL of headspace) which was composed of 90% of substrate at about 20 g COD L^−1^ (synthetic vinasse or aqueous solution of glucose) and 10% of inoculum at 2 g Volatile Suspended Solids (VSS) L^−1^ (*C. acetobutylicum* or *C. beijerinckii).* No nutritional supplementation was used. A 2 g NaHCO_3_ L^−1^ buffer solution was added to reaction medium in order to keep pH inside 4.0-6.0 interval throughout tests. Batch reactors containing synthetic vinasse plus *C. acetobulycum* and *C. beijerinckii* microorganisms were referred as VA and VB, respectively. According to the same nomenclature logics, batch reactors prepared with aqueous glucose solution plus *C. acetobulycum* and *C. beijerinckii* were labelled GA and GB.

### 2.4 Analytical methods and evaluation of batch reactors performance

Analyses of samples extracted from batch reactors were realized every 6 h for 96 h (total incubation time). Parameters measured (and their respective analytical technique for quantification) were the following: pH (4500-H+ electrometric method), COD (5220-D colorimetric method) (APHA, 2012), microorganism growth via OD (turbidity) (BERGOT *et al.*, 1996); concentration of carbohydrates into sucrose for batch reactors with synthetic vinasse as substrate (colorimetric method) (DUBOIS *et al.*, 1956); concentrations of glucose for batch reactors with aqueous glucose solution as substrate, SCOAs and alcohols (High Performance Liquid Chromatography – HPLC, Shimadzu Scientific Instrumentation®) (PENTEADO *et al.*, 2013).

HPLC facility was equipped with a degasifier (DGU-20A^3R^), two parallel pumps (LC-20AD), automatic sampler (SIL-20A_HT_), oven for heating up column (CTO-20^a^), ultraviolet detector with a diode arrangement (UV-DAD, SDP-20^a^) executing lectures at λ = 210 nm (wavelength), detector of refraction index (RID-10^a^) and controller (CBM-20A). Runnings were carried out in an Aminex

HPX-87H (300 mm x 7.8 mm, BioRad®) column at a volumetric flow rate of 0.5 mL min^−1^. Amount of injected volume of pre-filtered samples (its pH was previously adjusted to near 2.0) in chromatography apparatus was 100 μL. Furthermore, identification of peaks and calculation of concentrations of chemical species detected in samples through estimation of areas in chromatograms were assisted by LC Solution Shimadzu® software.

Performance of batch reactors in terms of some metabolites of interest was assessed by means of reaction production metrics such as productivity (*P,* mg L^−1^ h^−1^) and yield (*Y*, mg g^−1^), as defined in Eqs. (1) and (2), where *C_met,t_* (g met L^−1^) is the metabolite concentration at incubation time *t* (h); *C_met,t_*=0 is the initial concentration of metabolite at time *t* = 0; *C_i_* is the initial concentration of sugar (carbohydrates into sucrose or glucose).

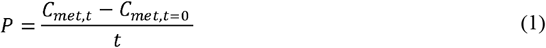

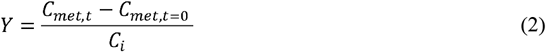

Mass balances in terms of theoretical COD were computed in order to check the compatibility between theoretical and experimental results at the end of batch experiments (incubation time of 72 h). For this purpose, atomic mass balance equations for each chemical element participating in total oxidation reaction of 1.0 mol of a organic compound (with general molecular formula C_x_H_y_O_z_, representing metabolites, carbohydrates and microbial biomass) were written to from stoichiometry provided by general reaction given in Eq. (3).

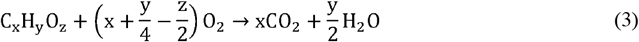

Therefore, since theoretical COD of a specific metabolite (*COD_t,met_*) corresponds to mass of oxygen required per unit of mass of organic compound oxidated, its expression is the one given in Eq. (4), where *x, y* and *z* are the numbers of carbon, hydrogen and oxygen atoms in organic molecule.

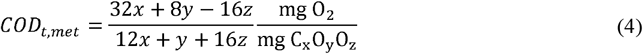

Total theoretical COD was calculated as a summation of COD weighted by concentrations of all batch process participants, namely carbohydrates, microbial biomass and metabolites, primarily short chain organic acids (SCOAs) and alcohols. It is noteworthy that carbohydrates (sucrose or glucose) and microbial biomass were also expressed as an organic compound with general molecular formula C_x_H_y_O_z_, so their respective theoretical CODs were determined through Eq. (4) as well. Particularly, for microbial biomass, specific molecular formula considered for *COD_t,BiO_* calculation was C5H9O3N (HONGLEI *et al.*, 2013), while *C_Bio_* was estimated from SSV content.

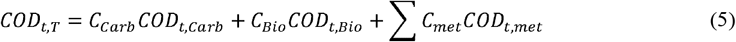

Aiming to check the consistency of total theoretical and experimental CODs (*COD_t,T_* and *COD_exp_,* respectively), a parameter named agreement (*Ag,* %), defined according to Eq. (7), was computed. So, the higher the value of *Ag*, the higher the consistency between *COD_t,T_* and *COD_exp_,* indicating that most of species generating during batch experiments was certainly detected experimentally.

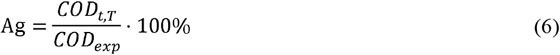

### 2.5 Kinetic modelling

Profiles of some selected participants of process (namely carbohydrates, lactic acid and microbial biomass) in batch reactors as a function of incubation time were adjusted by kinetic models. Main purpose of this curve fitting was to obtain reliable and simple equations to describe consumption and generation kinetics, which may be useful in process design, providing data for a confident computation of reactor scale up (LEVENSPIEL, 2000). Therefore, for consumption of carbohydrates (mono or disaccharide showing a decreasing profile of *C_carb_* (g Carb L^−1^) vs. t), process kinetic rate (*r_Carb_,* g Carb L^−1^ h^−1^) was modelled by classical first and second order equations, both defined according to the ordinary differential equation of Eq. (7), where *k* (adjustable) is the kinetic constant rate (units of h^−1^ for the first order reaction and L (g Carb^−1^) h^−1^ for second order) and *n* (fixed) is the reaction order (n = 1 for first order and *n* = 2 for second order) (ATKINS & DE PAULA, 2006).

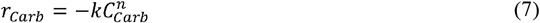

Kinetics of microorganisms’ growth was modelled via the original Gompertz equation as given by Eq. (8) with a little adaptation, which consisted simply in using *OD* (parameter used for monitoring cell multiplication in present work) instead of the logarithm of number of colony forming units (CFU). So, original Gompertz equation as in Eq. (8) was applied. Its adjustable parameters are the following: *C* is an empirical constant with the same units as *OD* (abs); *A* is a preexponential factor (abs); *e* is the Euler’s number (2,7182); *B* is the constant rate of maximum growing (h^−1^); t is the incubation time (h); *M* is the incubation time required to attain maximum growing (h) (GIBSON *et al.*, 1987; MCKELLAR & LU, 2004; BRUCKNER *et al.*, 2013).

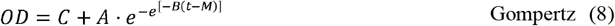

Additionally, *μ* (h^−1^) and *λ* (h), calculated by means of to Eqs. (9) and (10), are the specific growing rate and lag phase duration, respectively (BRUCKNER *et al.*, 2013).

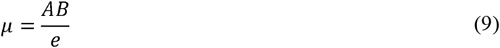

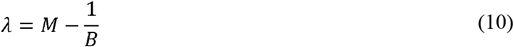

For some metabolites concentration profiles, modified Gompertz equation, typically employed to model accumulated hydrogen content in anaerobic digestion aiming biogas generation (CHEN *et al.*, 2006; AMORIM *et al.*, 2017), was used to describe kinetics of production.

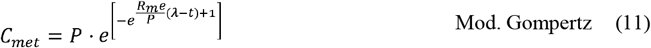

All kinetic models were fitted to experimental data using Levenberg-Marquadt optimization algorithm available in OriginPro 9.1® (ORIGINLAB, 2017) aiming to minimize the deviation between calculated and experimental values. Quality of nonlinear fitted curves were evaluated by means of two main parameters, namely the Residual Sum of Squares of (*RSS*) and the Coefficient of Determination (*R*^2^), which were given by OriginPro 9.1® right after the optimization procedure has been completed at the expense of a low number of iterations. They are defined by Eqs. (12) and (13), where *y_i_* is the experimental value at time *i*, *f_i_* is the correspondent fitted value at time *i* and 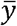 is the average of the experimental values set.

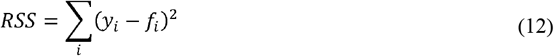

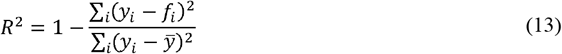

## 3. RESULTS & DISCUSSION

### 3.1 Removal of COD

Using synthetic vinasse prepared at 22.4 and 22.7 g COD L^−1^ (initial values of COD_sol_ near to 22 g COD L^−1^, see Table 1), reactors VA and VB showed average removal efficiencies of 3.6% and 4.8%, respectively. Final CODs reached in these media were of approximately 21.6 g COD L^−1^ after 96 h of incubation time. For experiments with glucose as substrate, 22.4 and 22.5 g COD L^−1^ were the starting CODs in reactors GA and GB. At the end of process (96 h), average removal efficiencies verified were 10.3% and 10.7% in reactors GA and GB, respectively, and both media attained a final COD of nearly 20.1 g COD L^−1^. This evidence of low degradation of COD can be imputed to the conversion of high amounts of carbohydrates existing in synthetic vinasse and aqueous glucose solution into metabolites, namely short chain organic acids (SCOAs) and ethanol, which account for increasing COD of reactional media, resulting in not very different initial and final CODs.

### 3.2 Profiles of cell growth, pH and carbohydrate content

Profiles of bacteria growth (OD), pH and total carbohydrate content for all batch reactors are presented in Fig. 1. Note that lines are plotted together with experimental points of pH profiles only with the purpose of guiding reader’s eye, while lines for bacteria cell growth and carbohydrate content profiles represent kinetic models fitted to experimental points of these monitoring parameters for experiments at different initial COD and pH.

**Fig. 1.**
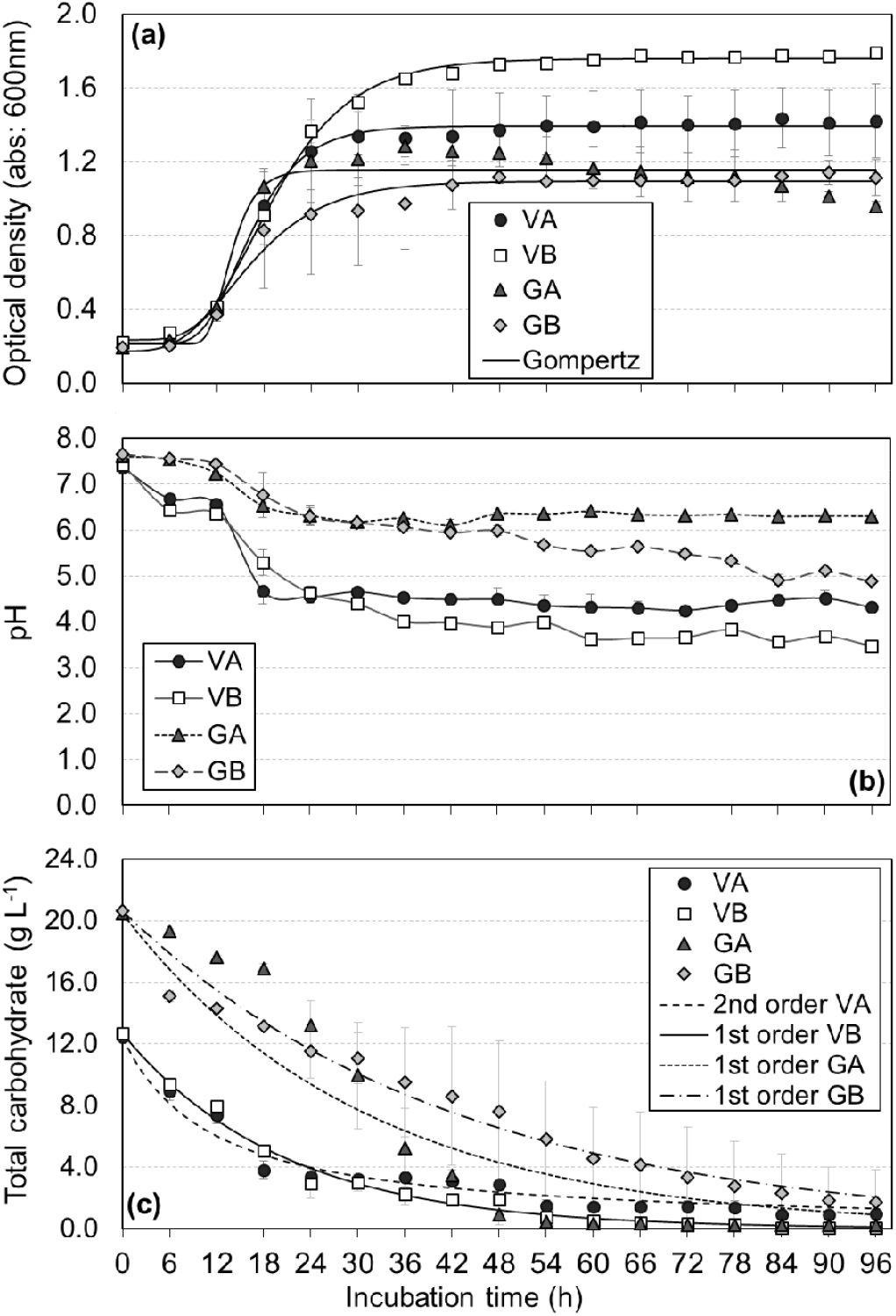
Profiles of some monitoring parameters in batch reactors along 96 h of operation: (a) cell growth by optical density (OD); (b) pH; (c) total carbohydrate content. Fitted kinetic models are plotted as lines for cell growth and total carbohydrate, while lines plotted for pH are just to guide the eye.

As shown Fig. 1a, growing trends of both microorganisms (*C. acetobutylicum* and *C beijerinckii,* measured by OD) were similar until 12 h in four experiments. However, most remarkable distinction among growing curves was observed from 24 h of incubation time, when reactor VB exhibited a greater number of cells until the end of process compared to other reactors. Additionally, reactor VA showed a higher stability of amount of microbial biomass after 30 h, while the same was verified for reactor VB solely after 48 h, for reactor GA in period 24-60 h and for reactor GB after 48 h. Cell multiplication in reactors containing synthetic vinasse was more expressive probably due to nutrients existing in this substrate (and not in aqueous glucose solution) have favoured growth of microorganisms.

Parameters obtained by fitting Gompertz model to experimental points of cell growth as expressed by OD are given in Table 2. Note that specific growth rates (*μ*) are higher for reactors containing *C. acetobutylicum* as inoculum (GA and VA), which also presented the longest lag phase periods (λ), corresponding to initial range of incubation time where no bacteria multiplication is observed. Also note that relatively low values for *RSS* (< 0.20) combined to relatively high values for *R^2^* (>0.90) are such that fitted Gompertz model (see solid lines in Fig. 1a) can be taken as really good to describe kinetics of increasing of microbial biomass in fermentation media.

**Table 2.** Model parameters, RSS and R^2^ for Gompertz model fitted to experimental points of cell growth (OD).

| Batch reactor | Fitted model parameters | | | | $\mu$<br>(h<sup>-1</sup>) | $\lambda$<br>(h) | $RSS$<br>(abs<sup>2</sup>) | $R^2$ |
| --- | --- | --- | --- | --- | --- | --- | --- | --- |
| | $C$ (abs) | $A$ (–) | $B$ (h<sup>-1</sup>) | $M$ (h) | | | | |
| VA | 0.21 | 1.18 | 0.22 | 14.60 | 0.10 | 10.08 | 0.01 | 0.9958 |
| VB | 0.23 | 1.53 | 0.15 | 16.67 | 0.08 | 9.87 | 0.01 | 0.9976 |
| GA | 0.21 | 0.94 | 0.47 | 12.96 | 0.16 | 10.82 | 0.12 | 0.9292 |
| GB | 0.17 | 0.92 | 0.15 | 13.66 | 0.05 | 7.14 | 0.03 | 0.9754 |

Regarding pH profiles shown in Fig. 1b, average initial value was ~7.5 for all batch reactors. Reactor VA presented pH stability in interval 4.0-4.5 from incubation time 18 h. Simultaneously, reactor VB exhibited a noticeable decreasing of pH from 0 h to 36 h, when a higher stability was attained, and pH oscillations were limited to range 3.5-4.0. Great reductions of pH in reactors VA and VB can be linked to a considerable production of acids in both reactional media, which will be showed and discussed latter in the present work. Moreover, after 30 h, reactor GA reached pH stabilization between 6.0-6.5, while in reactor GB the same stabilization was observed further ahead, just from 84 h, and pH values were kept around 5.0.

Evolution of total carbohydrate content in each one of fermentation media, which is depicted in Fig. 1c, indicates there was a high consumption by bacteria since remaining concentrations of both carbon sources were too low at the end of experiments. For reactors VA and VB, initial and final concentrations of carbohydrates were 12.5 and 0.9 g Carb L^−1^ and 12.6 and 0.2 g Carb L^−1^, implying carbohydrate conversion efficiencies of 92% and 98%, respectively. For reactors GA and GB, initial and final values were 20.5 and 0.2 g Glu L^−1^ and 20.6 and 1.7 g Glu L^−1^, associated to as high conversion efficiencies as those for media containing synthetic vinasse, of 99% and 92%. These results indicate in advance that increases of SCOAs and alcohols (two of the main classes of metabolites) concentrations in batch reactors should be pronounced. Nonetheless, this fact will be demonstrated lately here.

Parameters concerning first and second order kinetic models fitted to experimental points of carbohydrate content are presented in Table 3. For each one of substrates taken individually, results show that consumption of carbohydrate occurred faster in reactors VB and GA because their rate constants were the greatest ones, pointing out that, depending on substrate, *C. acetobulycum* (for synthetic vinasse) or *C. beijerinckii* (for aqueous glucose solution) species can exhaust carbon source more sharply over incubation time. Furthermore, based on *RSS* and *R^2^* values, one can conclude that best fitted models were precisely those plotted as dashed and solid lines in Fig. 1c. Therefore, first order model was the most suitable for describing profiles of carbohydrate content for reactors VB, GA and GB. For the same reason, second order model can be considered as the fittest for profile of carbohydrate content of reactor VA.

**Table 3.**
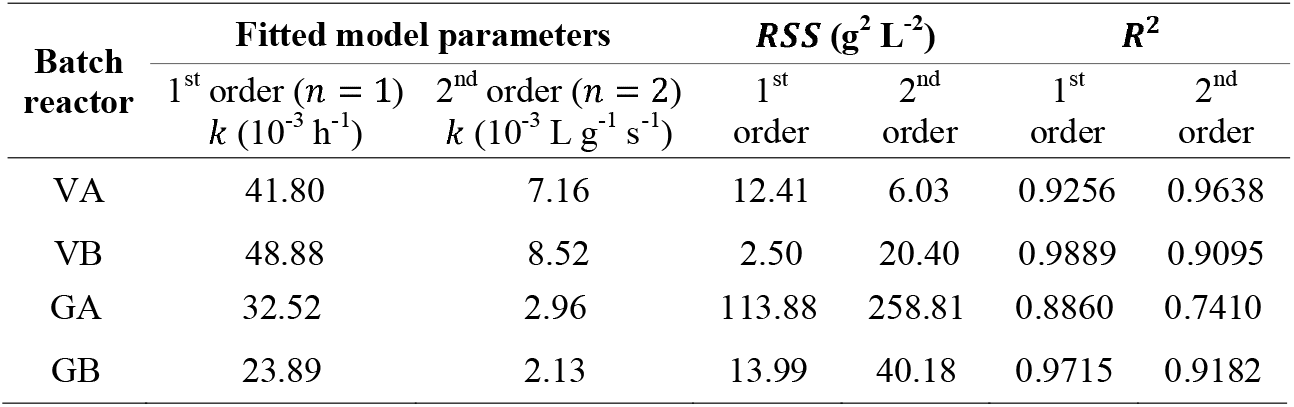
Model parameters, RSS and R^2^ for first and second order kinetic model fitted to experimental points of carbohydrate content.

| Batch reactor | Fitted model parameters | | $RSS$ (g<sup>2</sup> L<sup>-2</sup>) | | $R^2$ | |
| --- | --- | --- | --- | --- | --- | --- |
| | 1<sup>st</sup> order ($n = 1$)<br>$k$ (10<sup>-3</sup> h<sup>-1</sup>) | 2<sup>nd</sup> order ($n = 2$)<br>$k$ (10<sup>-3</sup> L g<sup>-1</sup> s<sup>-1</sup>) | 1<sup>st</sup> order | 2<sup>nd</sup> order | 1<sup>st</sup> order | 2<sup>nd</sup> order |
| VA | 41.80 | 7.16 | 12.41 | 6.03 | 0.9256 | 0.9638 |
| VB | 48.88 | 8.52 | 2.50 | 20.40 | 0.9889 | 0.9095 |
| GA | 32.52 | 2.96 | 113.88 | 258.81 | 0.8860 | 0.7410 |
| GB | 23.89 | 2.13 | 13.99 | 40.18 | 0.9715 | 0.9182 |

### 3.3 Concentration profiles of total SCOAs, lactic acid and ethanol

Concentration profiles of total short chain organic acids (T-SCOAs), lactic acid (lactate, HLa) and ethanol (EtOH) are depicted in Fig. 2. Process was shown to be strongly based on acidogenic fermentation with prevalence of lactic acid, but also generating other acids at considerably lower concentrations though. Particularly in reactors GA and GB, solventogenic fermentation resulting in more relevant generation of ethanol has occurred. Concentration of T-SCOAs accounts for concentrations of all acidic products detected during process using HPLC method. Acetic, butyric, propionic, isobutyric and malic acids were produced at appreciable amounts in batch reactors, however in a remarkably lower extent compared to lactic acid, which was indeed the major SCOA. Besides these metabolites, citric, succinic, formic, isovaleric, valeric and caproic were present in experiments at some incubation times and also contributed, even if not too significantly, for concentration of T-SCOAs.

**Fig. 2.**
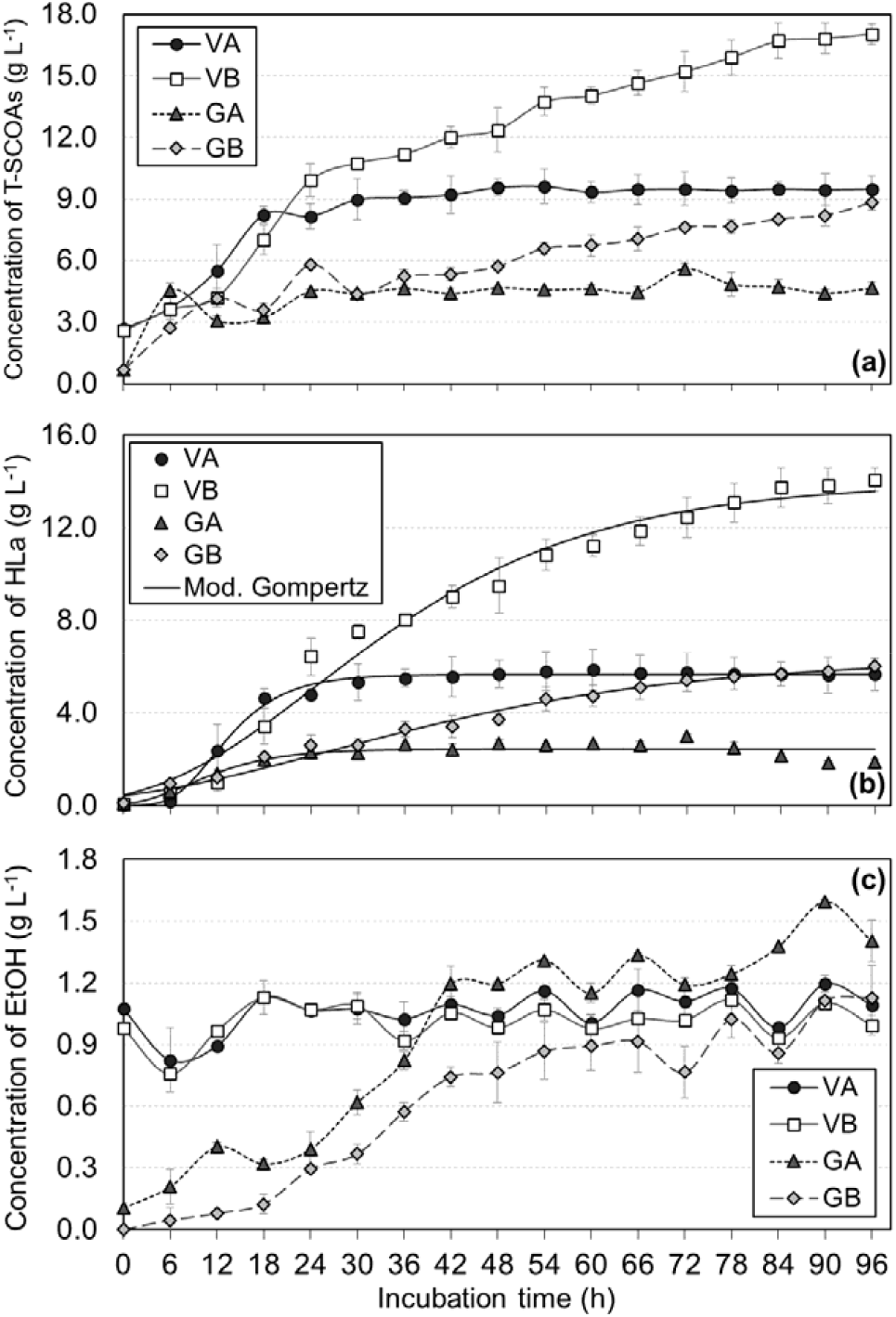
Profiles of metabolites concentrations in batch reactors along 96 h of operation: (a) total short chain organic acids (T-SCOAs); (b) lactic acid (HLa); (c) ethanol (EtOH). Fitted kinetic model is plotted as lines for lactic acid, while lines plotted for T-SCOAs and ethanol are just to guide the eye.

**Fig. 3.**
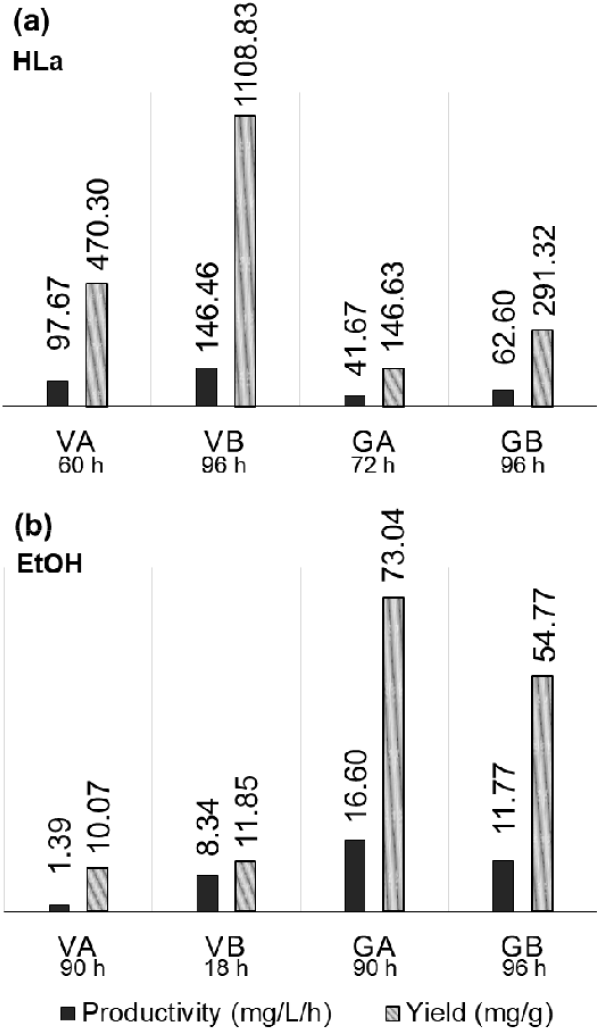
Productivities and yields of batch reactors at incubation times where the greatest concentrations of (a) lactic acid and (b) ethanol were verified.

Both in reactors VA and VB, concentration of T-SCOAs was initially of 2.6 g T-SCOAs L^−1^ because prepared synthetic vinasse already contained predefined amounts of butyric, acetic and propionic acids, see Fig. 2a. After 30 h of incubation time, T-SCOAs content in reactor VA remained nearly constant, while reactor VB showed a continuous increasing of it. As a consequence, concentrations were 9.5 g T-SCOAs L^−1^ in reactor VA and 17.0 g T-SCOAs L^−1^ in reactor VB at 96 h. Total acids concentration peaks were 5.6 g T-SCOAs L^−1^ for reactor GA at 72 h and 8.8 g T-SCOAs L^−1^ for reactor GB at 96 h, see Fig. 2a again. After 24 h has passed, concentration of T-SCOAs attained stability in reactor GA, while it was kept increasing continuously in reactor GB. So, these results point out to a likely productive advantage of *C. beijerinckii* compared to *C. acetobutylicum* in acidogenic fermentation under conditions used in this work for experiments using synthetic vinasse and aqueous glucose solution.

By observing Fig. 1a and Fig. 2a together, one can reasonably assume that pH and T-SCOAs concentration profiles seems to be tied to each other since acids content is determining for pH of fermentative media. Both parameters presented lesser variations after incubation time of 30 h in reactor GA. As expected in reactor GB, behaviours of pH and T-SCOAs were also opposite, although pH almost ceased decreasing after 18 h and T-SCOAs concentrations continued to increase sharply, reaching its maximum of 14.1 g HLa L^−1^ at 96 h, which was the greatest lactate content amongst those measured for all batch reactors. These same correlations are applicable to lactic acid in both GA and GB reactors, as can be checked by analyzing Fig. 1a and Fig. 2b.

Model parameters computed by fitting Gompertz modified model to experimental points of lactic acid are shown in Table 4. It is remarkable that reactor VB exhibited the greatest production potential of lactate (), which is compatible to the fact that it showed the highest concentration of lactic acid (Fig. 2 b). Reactor VA was the one showing the most pronounced lactic acid production rate (), which is certainly due to the steeper slope of its lactate profile in period just right after lag phase compared to those of other lactate profiles. Additionally, reactor VA also presented the longest lag phase time duration (), while it was zero for reactor GB since its lactate profile is the only one where it cannot be seen a clear distinguishable initial period where lactic acid content was nearly constant before experiments a pronounced increase.

**Table 4.** Model parameters, RSS and R^2^ for Gompertz modified model fitted to experimental points of lactic acid concentration.

| Batch reactor | Fitted model parameters | | | RSS<br>(g<sup>2</sup> L<sup>-2</sup>) | $R^2$ |
| --- | --- | --- | --- | --- | --- |
| | $P$ (g L<sup>-1</sup>) | $R_m$ (10<sup>-2</sup> g L<sup>-1</sup> h<sup>-1</sup>) | $\lambda$ (h) | | |
| VA | 5.65 | 41.50 | 6.31 | 0.42 | 0.9919 |
| VB | 13.95 | 25.81 | 4.64 | 6.75 | 0.9792 |
| GA | 2.44 | 14.36 | 1.98 | 1.32 | 0.8407 |
| GB | 6.35 | 8.92 | 0.00 | 0.85 | 0.9827 |

**Table 5.** Concentrations of the main metabolites obtained in batch fermentation tests using different substrates and pure cultures of Clostridium spp.

| Substrate | *Clostridium* strain | Concentrations of SCOAs and solvents (g L<sup>-1</sup>) |  |  |  |  |  |  | Reference |
| --- | --- | --- | --- | --- | --- | --- | --- | --- | --- |
|  |  | HBu | HAc | HLa | HPr | AcO<sup>#</sup> | BuOH | EtOH |  |
| Glucose<br>(20 g Glu L<sup>-1</sup>) | *C. acetobutylicum* ATCC 824 | 3.87 | 2.40 | NA | NA | 0.02 | 0.05 | NA | Holt *et al.* (1984) |
| Glucose<br>(40 g Glu L<sup>-1</sup>) | *C. acetobutylicum* ATCC 824 | 4.96 | 4.92 | NA | NA | 0.41 | 1.33 | 0.28 | Holt *et al.* (1984) |
| Glucose<br>(10 g Glu L<sup>-1</sup>) | *C. acetobutylicum* ATCC 824 | 5.4 | 1.2 | NA<sup>+</sup> | NA | NA | 0.2 | 0.5 | Oh *et al.* (2009) |
| Sugar mix<br>(28 g Suc L<sup>-1</sup> + 4 g Glu L<sup>-1</sup> + 8 g Fru<sup>**</sup> L<sup>-1</sup>)<br>(48 h)<sup>§</sup> | *C. saccharobutylicum* DSM 13864 | 0.72 | 13.15 | NA | NA | 3.88 | 8.79 | 0.48 | Ni *et al.* (2012) |
| Cane molasses<br>(60 g total sugar L<sup>-1</sup>) (84 h) | *C. beijerinckii* L175 | ~1.2 | ~1.1 | NA | NA | ~4.8 | ~15.0 | ~2.2 | Li *et al.* (2013)* |
| Cheese whey<br>(55 g Lac<sup>e</sup> L<sup>-1</sup>) | *C. acetobutylicum* ATCC 824 | 1.70 | 2.61 | 5.49 | NA | 0.04 | 0.06 | 1.20 | Durán-Padilla *et al.*<br>(2014) |
| Glucose (40 g Glu L<sup>-1</sup>)<br>(43 h) | *C. saccharobutylicum* DSM 13864 | 0.15 | 0.45 | 0.20 | NA | 3.62 | 5.97 | 0.27 | Magalhães *et al.* (2018) |
| Xylose (40 g Xyl<sup>f</sup> L<sup>-1</sup>)<br>(43 h) | *C. saccharobutylicum* DSM 13864 | 0.57 | 0.90 | 0.32 | NA | 2.32 | 4.35 | 0.24 | Magalhães *et al.* (2018) |
| Glucose (80 g Glu L<sup>-1</sup>)<br>(72 h) | *C. acetobutylicum* ATCC 824 | 0.22 | NA | 2.33 | NA | NA | NA | 2.21 | Guilherme *et al.* (2018) |
| Synthetic vinasse<br>(20 g DQO L<sup>-1</sup>; 12 g Suc L<sup>-1</sup>)<br>(96 h) | *C. acetobutylicum* ATCC 824 | 0.72 | 1.64 | 5.65 | 0.67 | NA | NA | 1.09 | Present work |
| Synthetic vinasse<br>(20 g DQO L<sup>-1</sup>; 12 g Suc L<sup>-1</sup>)<br>(96 h) | *C. beijerinckii* ATCC 25752 | 0.57 | 0.96 | 14.06 | 0.89 | NA | NA | 0.99 | Present work |
| Glucose (20 g Glu L<sup>-1</sup>)<br>(96 h) | *C. acetobutylicum* ATCC 824 | 0.35 | 0.40 | 1.85 | 0.40 | NA | NA | 1.40 | Present work |
| Glucose (20 g Glu L<sup>-1</sup>)<br>(96 h) | *C. beijerinckii* ATCC 25752 | 0.65 | 0.61 | 6.01 | 0.54 | NA | NA | 1.13 | Present work |

<sup>#</sup>AcO: acetone; <sup>\*\*</sup>Fru: fructose; <sup>e</sup>Lac: lactose; <sup>f</sup>Xyl: xylose; <sup>+</sup>NA: not available or not detected; <sup>§</sup>Incubation time where data were collected were reported, so it was included here; \*Concentrations preceded by the symbol “~” were extracted from plots, so they are approximations.

It possible to assert with good level of certainty that trend of lactic acid production was determining for trend of T-SCOAs production even because lactate was the major metabolite detected in all batch reactors of present work. Thus, in the same way as for lactic acid concentration, an uninterrupted increase of T-SCOAs concentration was observed in reactor VB from incubation time of 30 h until the end of experiment. In parallel, a greater stability was verified for both T-SCOAs and lactic acid concentrations in reactor VA, showing variations limited to 5.0-6.0 g HLa L^−1^ of lactate content. For reactor GB, the highest lactate content, 6.0 g HLa L^−1^, was attained at the end of process, whereas for reactor GB lactate concentration was kept in range 2.0-3.0 g HLa L^−1^ from 24 h. Based on these results, one can realize there is a direct relationship of cause and effect between increasing lactate concentration (cause) and unceasing decrease of pH of reactional media (effect).

Moreover, it is important to highlight that a similar trend to the ones revealed in Figs. 2a and 2b is also observed in acidogenic reactors intended to produce hydrogen (H_2_), where production of lactic acid is a strong indicative of reduction in generation of biogas and other acidic products since lactate generation depletes the carbon source which would otherwise be used by bacteria to provide these compounds as metabolites (MARTINS & AMORIM, 2016). As a consequence, lactate tends to be more decisive for T-SCOAs concentration profile. This is additionally corroborated by chemical reactions written in Eqs. (8)-(10) considering glucose as substrate.

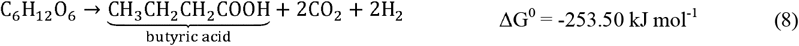

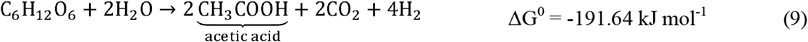

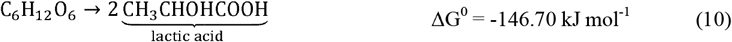

According to Jones & Woods (1986), *C. acetobutylicum* is able to convert pyruvate resulting from oxidation of initial carbohydrate content (HOELZLE *et al.*, 2014) into lactate under certain circumstances. At normal conditions, however, lactic acid route is not the preferred one from energy perspective since its ATP generation is lower than that of butyrate and acetate routes, see Eqs. (8) and (10), although it seems to be a less efficient alternative to allow energy generation and oxidation of NADH to continue occurring when mechanisms of making protons and electrons available through production of H_2_ are blocked (JONES and WOODS, 1986).

Despite of frequently receiving less attention than butyric and acetic acids, lactic acid is also a metabolite that can be obtained during ABE fermentation. Lactate is not taken as a precursor for solvents production such as acetate and butyrate are, but it is considered as a secondary compound resulting from a deviation in the production route of butanol (LÜTKE-EVERSLOH & BAHL, 2011). Concerning it, according to Bahl *et al.* (1986) and Durán-Padilla *et al.* (2014), there is a greater propensity to produce more lactic acid in an iron-deficient fermentation medium maintained at pH higher than 5.0. Under these conditions, synthesis of chemical intermediates and activity of enzymes involved in ABE fermentation, such as ferredoxin and pyruvate ferredoxin oxidoreductase, are harmed, which leads to redirection of metabolic flow to lactate (DURÁN-PADILLA *et al.*, 2014).

According to Hoelzle *et al.* (2014), capacity to produce lactic acid is very common amongst microorganisms, although this chemical is generally not considered as a primary metabolite except if fermentative microbial is of *Lactobacillus* genus. In this sense, for many microbial species, lactate serves as an intermediate in anaerobic process, being obtained in an earlier stage to disappear right subsequently in process (HOELZLE *et al.*, 2014). Notwithstanding, one cannot observe periods of lactate consumption in profiles of both reactors VB and GB in Fig. 2b, pointing out that this metabolite was not obtained as a chemical intermediate, but most probable as a final product notably in experiments performed with *C. beijerinckii* in this work.

Durán-Padilla *et al.* (2014) have performed anaerobic digestion experiments of cheese whey (initial lactose content of 55 g Lac L^−1^) without iron supplying, which resulted in fermented media containing lactic acid as the major metabolite. Comparing these to results of present work, one cannot surely assert that iron amount used for preparing synthetic vinasse (10 mg Fe^3+^ L^−1^, see Table 1) was scarce, although this is a hypothesis deserving further investigation. Moreover, Oshiro *et al.* (2010) studied the influence of previous addition of lactic acid to reactional media of ABE batch fermentation and verified an inhibition of glucose and lactate (latter also can serve as carbon source) consumption at 10 and 20 g HLa L^−1^, which was associated to acidic stress. So, taking results of reactors GA and GB into analysis points out there may have been a restriction of *Clostridium* metabolism due to high contents of lactate, preventing butanol generation. In the same sense, Zhou *et al.* (2018) concluded that excessive supplementation of substrate with lactic acid (20 g Glu L^−1^ + 5 g HLa L^−1^ or more) is lethal to *C. saccharoperbutylacetonicum* N1-4 (ATCC 27021), making solvent production unfeasible. However, no clear evidence of mortality of microbes’ cells was observed in batch reactors of this work, so the deviation of carbon source to production of lactate instead of butanol is more presumable.

As shown in Fig. 2c, initial ethanol concentrations were 1.07 g EtOH L^−1^ in reactor VA and 0.98 g EtOH L^−1^ in reactor VB, both consistent with ethanol content defined for synthetic vinasse (see Table 1). For first 6 h, ethanol concentrations were reduced to 0.83 and 0.76 g EtOH L^−1^ in reactors VA and VB, respectively. During period of 6-18 h there were slight decreases in both reactional media, while in 18-96 h it was observed a greater stability within of 1.0-1.2 g EtOH L^−1^ and 0.9-1.2 g EtOH L^−1^ in reactors VA and VB, respectively. Based on these results, one can claim there was no appreciable production of ethanol in batch reactors using synthetic vinasse, so solventogenesis did not occur in view that other solvent were also not generated. Decreasing of pH to 4.5 or lower values in reactors VA and VB (see Fig. 1b) is a signal that substrate was mainly directed to production of SCOAs, resulting in a substantial decrease of pH, although this could also favour solvent generation (LI *et al.*, 2014)

Both batch reactors using glucose as substrate (GA and GB) showed considerable ethanol production over some incubation periods, but also experimenting periods of ethanol consumption of lesser magnitude. Peaks of average ethanol concentration were 1.60 g EtOH L^−1^ in reactor GA at 90 h and 1.13 g EtOH L^−1^ in reactor GB at 96 h. In this sense, according to Haus *et al.* (2011), it is possible that ethanol generation occurs during acidogenesis independently of production of other solvents in ABE fermentation with *C. acetobutylicum,* which can serve to explain increasing profiles of reactors GA and GB given in Fig. 2c since other solvents except ethanol were not detected in the present work.

Absence of butanol in reactors GA and GB can be linked to low glucose content in substrate, which was 20 g Glu L^−1^ and is definitely below those of previous works where butanol was detected LI *et al.*, 2011; NI *et al.*, 2012; GUILHERME *et al.*, 2018). As a consequence, production of SCOAs was prevalent over solvents not only in experiments with glucose of this work, but also in those using synthetic vinasse. In fact, acidogenic fermentation tends to be the prevailing one especially at low concentrations of sugars in substrate (MONOT *et al.*, 1982; MAGALHÃES *et al.*, 2018). Additionally, it is noteworthy that even at 60 g Gli L^−1^ of initial concentration of carbon source, there are technical literature reports on cases of poor butanol generation (3.7 mg BuOH L^−1^) in a fermentation medium under pH controlled at 6.8 and 5.0 (GEORGE & CHEN, 1983).

For lactic acid and ethanol, productivities and yields computed as defined by Eqs. (1) and (2) are shown Fig. 5, where one can clearly observe that best production metrics are those associated to reactor VB for lactate and reactor GA for ethanol, which occurred at 96 h and 90 *h*, respectively. Therefore, without losing sight of conditions defined for carrying out experiments, *C. acetobulycum* ATCC 824 provided higher performance parameters for lactate in batch reactors using synthetic vinasse, while the same was verified for *C. beijerinckii* ATCC 25752 in batch reactors using aqueous glucose solution.

### 3.4 Concentration profiles of other relevant SCOAs obtained by acidogenic fermentation

Profiles of some others SCOAs quantified during experiments, namely butyric, acetic and propionic acids (butyrate, HBu; acetate; HAc; propionate, HPr), are presented in Fig. 4. Batch rectors using synthetic vinasse were operated beginning from predefined concentrations of butyric, acetic and propionic acids (see Table 1). Butyric acid profile (Fig. 4a) started from 0.53 g HBu L^−1^ in reactors VA and VB. For reactor VA, maximum of butyrate concentration was near 1.0 g HBu L^−1^ at 6 h followed by a decrease until 12 h and finally a stability period of 12-96 h in range 0.7-0.8 g HBu L^−1^. Reactor VB showed a peak of approximately 0.8 g HBu L^−1^ at 12 h and subsequent period was of stability between 0.5-0.6 g HBu L^−1^. Initial concentrations of acetate were 1.30 and 1.17 g HAc L^−1^ in reactors VA and VB (Fig. 4b), respectively, where maximum of 1.64 and 1.37 g HAc L^−1^ were detected at 18 h and 42 h. Level of acetate was practically constant at 1.64 g HAc L^−1^ in period 18-96 h in reactor VA, while in reactor VB it was reduced to 0.95 g HAc L^−1^ at 60 h and after that it was kept under low magnitude oscillations until 96 h. Additionally, for both reactors VA and VB, initial concentrations of propionic acid were nearly 0.55 g HPr L^−1^ (Fig. 5c). The highest lactate profile was the one concerning reactor VB, where concentration showed a peak of 1.03 g HPr L^−1^ at 18 h and subsequently remained in interval 0.8-0.9 HPr L^−1^. For reactor VA, propionate content exhibited a maximum of 0.85 g HPr L^−1^ and after 24 h it was limited to 0.6-0.7 g HPr L^−1^.

**Fig. 4.**
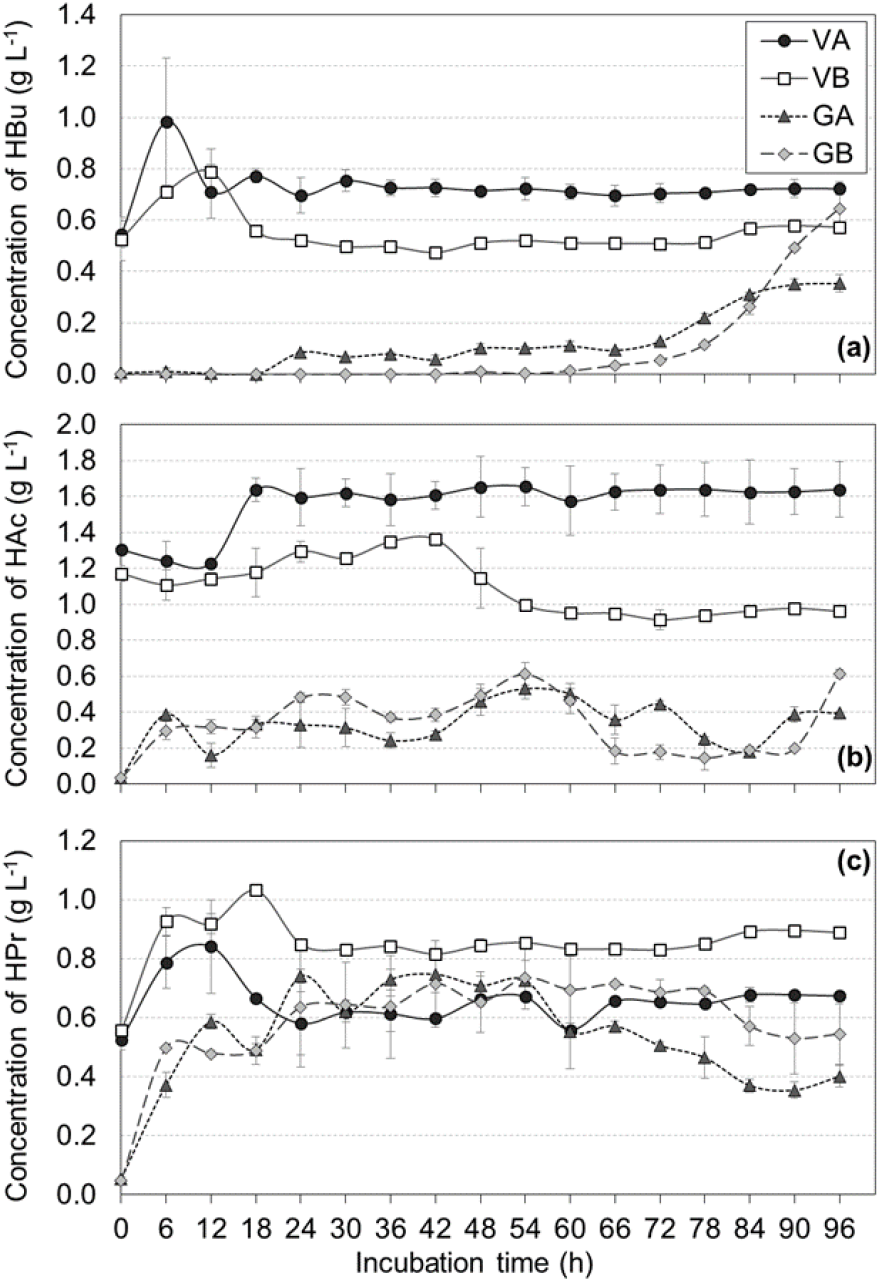
Profiles of some metabolites concentration in batch reactors along 96 h of operation: (a) butyric acid (HBu); (**b)** acetic acid (HAc); (c) propionic acid (HPr). Lines are plotted just to guide the eye.

In reactors using glucose as substrate, butyrate concentration presented maximum at the end of experiments, which were 0.36 and 0.65 g HBu L^−1^ in reactors GA and GB, respectively. Lactate was produced and consumed along alternating periods in batch reactors GA and GB, where its highest concentrations were 0.39 and 0.47 g HAc L^−1^, both at 96 h. Moreover, reactor GB showed acetic acid contents mostly greater than those of reactor GA. For propionate concentration, which did not show wide variations during experiments with glucose, reactor GA exhibited a maximum of 0.74 g HPr L^−1^ at 54 h, while it was 0.73 g HPr L^−1^ at 54 h in reactor GB.

Excepting malic acid in reactors GA and GB, which presented concentration maximum of 2.35 and 1.53 g HMa L^−1^ at 6 h and 12 h, respectively, other acidic metabolites were quantified at notably lower contents during experiments. Succinic, isobutyric, isovaleric and valeric acids were detected in all batch reactors, while citric and formic acids were present only in reactors GA and GB and caproic acid only in reactor VA. Moreover, methanol and butanol also were not quantified at no incubation times.

Butyric and acetic acids are both precursors of solvents production through ABE fermentation process. These acidic metabolites resulting from acidogenesis stage of biochemical route are required at certain concentrations in reactional medium to trigger the solventogenesis stage. For *C. acetobutylicum* ATCC 824, Monot *et al.* (1982) estimated that butyrate content should be at least 1.9 g HBu L^−1^ in order to the onset for solvents production to be attained. Since neither in reactor VA nor in reactor VB showed even lower concentrations of butyrate over incubation time, so it can be linked to inhibition of solventogenesis in experiments with synthetic vinasse of this work. Hereupon addition of butyric and acetic acids, besides carbonates, acetate esters and others, has been investigated aiming to stimulate butanol generation by means of biochemical route and to improve performance of some bacteria strains (VENTURA & JAHNG, 2013; TSAI *et al.*, 2014; AL-SHORGANI *et al.*, 2018). Concentrations of precursors considered in all these previous works, however, were considerably higher than those defined for synthetic vinasse used in present work.

In this work, even though pure cultures of *C. acetobutylicum* and *C. beijerinckii* have been applied for fermentation of solutions of glucose and sucrose plus some supplements (synthetic vinasse), results indicated that concentrations of butyric and acetic acids were certainly not high enough to initiate butanol production because they were limited to maxima of 0.98 HBu L^−1^ and 1.64 g HAc L^−1^. Therefore, it is plausible to assert that higher concentrations of precursors acids would be required to provoke butanol generation, at least nearer those reported in technical literature some time ago as critical for *C. acetobutylicum* ATCC 824, such as 1.5 g HBu L^−1^ (MONOT *et al.*, 1984; HÜSEMANN e PAPOUTSAKIS, 1988) and 1.9 g HBu L^−1^ (MONOT *et al.*, 1982), for onsetting of solventogenesis.

Applying *C. acetobulycum* ATCC 824 to a substrate containing a number of supplements recommended by Gungorsmusler *et al.* (2010), at 20-80 g Gli L^−1^ of carbohydrate concentration and initial pH 6.5, Guilherme *et al.* (2018) have carried out batch ABE fermentation experiments intending to generate butanol, which was not obtained though. An exception was observed when agitation at 100 rpm was used (producing 0.41 g BuOH L^−1^), although this was not effective to induce solventogenesis in present work. Moreover, major metabolites detected were butyrate and acetate and carbon source depletion was too low, different from what was observed in present work, so this was taken as the cause for absence of solvents by Guilherme *et al.* (2018) since process followed through an essentially acidogenic pathway.

Li *et al.* (2011) solely detected solvents in batch ABE fermentation experiments after a certain period of incubation required for enough accumulation of butyric and acetic acids. Same behaviour was verified by George & Cheng (1983), Ventura *et al.* (2013) and Wang *et al.* (2014) for feed batch and batch processes. Previously, Holt *et al.* (1984) evaluated ABE fermentation by *C. acetobutylicum* ATCC 824 while keeping neutral pH for reactional media at 20 g Glu L^−1^, resulting in a predominantly acidogenic process with poor solvent generation (0.05 g BuOH L^−1^). However, addition of supplements such as butyrate, acetate and propionate was effective to induce a greater production of butanol, which corroborates the hypothesis that there was a deficit of acids precursors for solventogenesis during batch experiments of present work. Moreover, it also draws focus to pH control strategy of fermentation, which was not tested in this work, but certainly would imply a relevant difference (HOLT *et al.*, 1984). In this context, it is worth mentioning there are some bacteria strains known as low-acid-producing, such as *C. acetobutylicum* B18 (PARK *et al.*, 1993), which are able to perform solventogenesis from considerably low acids concentrations.

### 3.5 Summary of comparisons to previous works and results of mass balance

For comparison purposes, Table 2 summarizes some results from reference literature of batch experiments with pure *Clostridium* spp. cultures and a number of different substrates (some of these works were discussed earlier here) and those obtained in present work. Particularly, a closer quantitative accordance regarding lactic acid concentration is remarkable for batch test with synthetic vinasse (20 g DQO L^−1^) and that with cheese whey (55 g Lac L^−1^) by Durán-Padilla *et al.* (2014), both applying *C. acetobutylicum* ATCC 824, although differences related to other acidic metabolites and also solvents are outstanding. Moreover, one can realize that lactate concentrations obtained in experiments of this work are notably greater than those reported in other works listed in Table 2.

Additionally, mass balances computed with concentrations of total carbohydrate, metabolites and microbial biomass at incubation time 96 h for all batch reactors are found in Table 3. Agreement parameter varied from 50% (reactor GA) to 99% (reactor VB), indicating there was probably generation of liquid organic products not detected and quantified by means of HPLC method or even gases dissolved in media resulting from fermentation (LEE *et al.*, 2003; PEIXOTO *et al.*, 2012; MARTINS & AMORIM, 2016). This is particularly applicable for batch reactors VA, GA and GB, but more pronouncedly for reactor GA, where difference between COD_t,T_ and COD_exp_ was the greatest one. For an acidogenic process aiming to generate biogas (H_2_) from natural fermentation of coconut industrial wastewater, Martins & Amorim (2016) also computed mass balance based on COD and obtained values for agreement parameter comparable to those shown in Table 6 concerning results of this work.

**Table 6.**
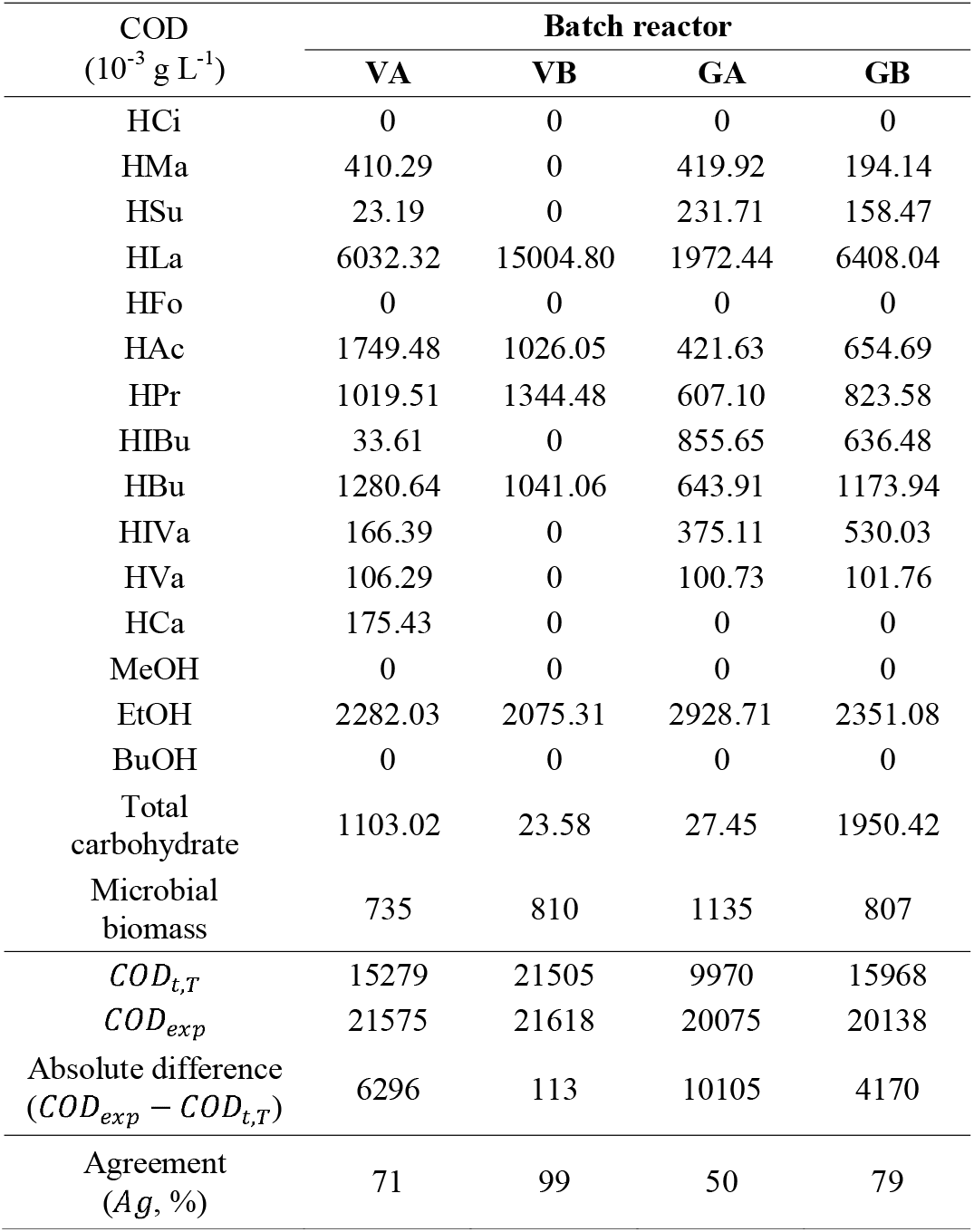
Mass balance in terms of COD at the end of operation (96 h) for all batch reactors.

## 4. CONCLUDING REMARKS

Butyric and acetic acids were detected at appreciable amounts, however not high enough to induce the onset for production of butanol, which was not detected at any incubation times along experiments at all. Therefore, distinctive profiles of complete ABE fermentation process, showing acidogenic stage followed by the solventogenic one, were not observed. Instead, profiles for batch reactors indicated that lactate production pathway seemed to be rather followed than others since lactic acid was the major metabolite.

Synthetic vinasse prepared at laboratory already presented concentrations of butyrate, acetate, propionate and ethanol, therefore generation of these metabolites in reactors VA and VB were poor because their concentrations were not very higher or sometimes lower than the initial ones. In contrast, production of them was more expressive in reactors GA and GB even because they did not exist in original substrate. Moreover, despite of ethanol have not been generated along experiments using synthetic vinasse, it was obtained in reactors containing glucose probably through a pathway unrelated to those for other solvents. This is the main distinction between results of experiments using synthetic vinasse and aqueous glucose solution of this work.

Chemicals present in synthetic vinasse, mainly inorganic salts and organic acids, may have had some influence on production of metabolites in both reactors VA and VB. Greater lactic acid contents were obtained in reactor VB. In batch reactors using glucose, concentrations of T-SCOAs and individual SCOAs were predominantly more expressive with *C. beijerinckii* (reactor GB), although ethanol generation has been more pronounced with *C. acetobutylicum* (reactor GA) in mostly of incubation times. Moreover, it is also highlightable that reactors containing glucose as substrate exhibited higher ethanol contents than those with synthetic vinasse.

Although it cannot be surely asserted, a probable cause for absence of butanol in reactional media was the low concentration of carbon source available in substrates, which was mostly directed to lactate production in all batch experiments according to metabolites profiles, showing that process was essentially an acidogenic fermentation with a more abundant generation of lactate instead of other typical products. So, in view of results of present work, it is presumable there was a premature interruption of biochemical production of butanol in reactors containing both selected microbes and substrates.

## ACKNOWLEDGMENTS

Authors are grateful to CNPq (Brazilian National Council for Scientific and Technological Development – Finance Code 142340/2016-2) and the São Paulo Research Foudation (FAPESP, BIOEN Program, grants number 2012/09785-8) for scholarship and financial support for research.

